# Biochemical properties of glycerol kinase from the hypersaline-adapted archaeon *Haloferax volcanii*

**DOI:** 10.1101/2025.02.14.638313

**Authors:** Karol M. Sanchez, Julie A. Maupin-Furlow

## Abstract

Extremophilic microorganisms hold promise to serve as robust biocatalysts in the conversion of glycerol waste into high value products. *Haloferax volcanii* is a hypersaline-adapted archaeon that prefers glycerol over glucose and channels this carbon source into central metabolism through glycerol kinase (GK). Here we report the biochemical properties of the *H. volcanii* GK and evaluated its potential for biotechnological applications. The N-terminal His-tagged form of GK was found functional *in vivo* and was readily purified to homogeneity at 4.5-fold higher yield (3 mg/L culture) than GK fused to a C-terminal StrepII tag. Further analysis of His-GK by size exclusion chromatography revealed the enzyme exhibited a glycerol-induced shift from a homodimer to a homodimer-homotetramer equilibrium. Purified His-GK demonstrated robust activity over a broad pH and salinity range, with optimal activity at 100 mM NaCl and 50-60 °C. The enzyme was catalytically active in organic solvent (5-10 % DMSO) and crude glycerol containing methanol. His-GK was also found to exhibit full activity after freeze-thaw, showed prolonged thermotolerance in 2 M NaCl supplemented buffers, and had a melting temperature (T_m_) in the range of 83-84 °C. Kinetic analysis using the Hill equation indicated His-GK displayed positive cooperativity for glycerol, ATP, and magnesium, with manganese and cobalt also found to serve as divalent cation cofactors. These findings underscore the unique and robust enzymatic properties of *H. volcanii* GK, representing the first known GK to exhibit positive cooperativity with glycerol and ATP, and highlighting its potential for biotechnological applications in glycerol conversion.

**IMPORTANCE:** This study provides new insight into glycerol kinase kinetics through the discovery of an enzyme exhibiting positive cooperativity for glycerol and ATP. By purifying and characterizing *H. volcanii* glycerol kinase, this work marks the first isolation of a glycerol kinase from a halophilic archaeon. The enzyme displays unique properties, including resilience to organic solvents, high temperatures, and extreme salinity. It also exhibits sigmoidal kinetics, with Hill coefficients averaging n = 2 for glycerol, ATP, and magnesium, indicating positive cooperativity. This behavior, previously unobserved in other glycerol kinases, aligns with the preference of *H. volcanii* for glycerol over glucose. Given the role of glycerol kinases in converting glycerol waste into value-added products, their efficiency is often a bottleneck in bacterial systems that prefer glucose and struggle under extreme conditions. The distinctive properties of *H. volcanii* glycerol kinase suggest potential for biotechnological applications in harsh environments.

## INTRODUCTION

Glycerol is an abundant carbon source with the potential to be used in numerous applications in the cosmetic, pharmaceutical, food, and renewable chemical industries (1). The biodiesel industry generates vast quantities of glycerol as a byproduct, with approximately one pound produced for every gallon of biodiesel (2). This excessive accumulation of glycerol presents significant industrial challenges due to its high volume, currently limited market applications, and risk of environmental pollution (2, 3). However, glycerol waste streams also provide opportunities to transform this three-carbon intermediate into valuable chemicals such as 1,3-propanediol, butanol, ethanol, biodiesel, biomethane, monoglycerides, citric acid, and docosahexaenoic acid (2, 4, 5). These transformations require robust microorganisms that display preference for glycerol as a carbon source and enzymes capable of functioning efficiently under industrial conditions (2, 4).

The hypersaline-adapted archaeon *Haloferax volcanii*, originally isolated from the Dead Sea (6), has garnered attention based on its robust growth in harsh conditions and preference for glycerol as its primary energy and carbon source (7–11). By contrast, most organisms use a ‘salt-out’ strategy that relies upon counterbalancing the osmotic stress through production of organic osmolytes, such as glycerol, which can leak out into the environment for use by the halophilic archaea (12). Thus, *H. volcanii* has unusual properties, not found in most organisms, including the preference for glycerol over glucose, and an entire proteome adapted with halophilic, organic solvent tolerant, and thermophilic properties including surfaces reduced in hydrophobicity and enriched acidic residues compared to ‘mesohalic’ counterparts (13–16). These adaptations make haloarchaeal enzymes attractive for industrial processes, which often involve extreme conditions. Moreover, *H. volcanii* transcription factors, GlpR, TrmB, TbsP and GfcR, are identified that control expression of central metabolic pathways providing additional tools to direct carbon flow through synthetic biology (17–20). The halophilic properties, ability to tolerate harsh growth conditions, and carbon source preferences enhance the suitability of this archaeon for industrial and biotechnological applications (8).

Glycerol kinase (GK) plays a central role in glycerol metabolism. In *H. volcanii*, glycerol is metabolized through the action of GK and glycerol-3-phosphate (G3P) dehydrogenase to generate 1,3-dihydroxyacetone phosphate (DHAP) for central metabolism (7, 21). Using ATP, GK phosphorylates intracellular glycerol to form G3P and ADP as a byproduct. This metabolic step is essential for glycerol consumption based on finding GK-deficient mutants are unable to grow on glycerol compared to other carbon sources (7). Optimizing the purification and determining the biocatalytic properties of haloarchaeal GKs are crucial for advancing fundamental knowledge and broadening the applications of these GKs in the bioindustry.

Here we report the purification and biochemical characterization of the first haloarchaeal GK and, thus, address a significant gap in understanding GK biochemistry. This study focused on *H. volcanii* GK, as it represents a promising candidate for the biotransformation of glycerol into value-added chemicals. We find the *H. volcanii* GK displays resilient properties and provide the first example of a GK that exhibits positive cooperativity for glycerol, ATP and magnesium. The *H. volcanii* GK is observed to undergo a glycerol-induced a conformational change from dimer to a tetramer consistent with its apparent positive allosteric control by glycerol. Thus, the *H. volcanii* GK is found to be fundamentally distinct from characterized GKs, expanding our understanding of extremophilic enzymes and their potential for glycerol bioconversion in industries like the biodiesel sector. As archaeal metabolism is advanced, research on *H. volcanii* adds to the growing arsenal of biotechnological tools and innovations.

## MATERIALS AND METHODS

### Strains and media

Strains and plasmids utilized in this study are listed in Table 1. *Escherichia coli* strain Top10 (Life Technologies, Carlsbad, CA) was used for routine cloning procedures. To obtain plasmids devoid of methylation for subsequent transformation into *Haloferax volcanii*, *E. coli* strain GM2163 (New England Biolabs, Ipswich, MA) was employed, following the protocol described by Cline *et al*. (22). Liquid cultures were aerated by orbital shaking at 200 rpm. *E. coli* strains were cultured at 37 °C in Luria-Bertani (LB) medium, with ampicillin (100 mg·L⁻¹) added when required. *H. volcanii* strains were grown at 42 °C in American Type Culture Collection 974 rich medium (ATCC974) or minimal medium supplemented with 20 mM glycerol (GlyMM) or fructose (FMM). Media was prepared according to the protocols in *The Halohandbook* (23). Growth media supplements included novobiocin (0.2 μg·mL⁻¹), 5- fluoroorotic acid (5-FOA, 50 μg·mL⁻¹), and uracil (50 μg·mL⁻¹). Before adding to the medium, 5-FOA was dissolved in 100 % dimethyl sulfoxide (DMSO) at 50 mg·mL⁻¹.

**Table 1.** List of strains and plasmids used in this study^a^.

| Strain or plasmid | Description | Source or reference |
| --- | --- | --- |
| ***E. coli* strains** |  |  |
| Top 10 | F– *recA1 endA1 hsdR17(r<sub>K</sub>–m<sub>K</sub>+) supE44 thi-1 gyrA relA1* | Invitrogen |
| GM2163 | F– *ara-14 leuB6 fhuA31 lacY1 tsx78 glnV44 galK2 galT22 mcrA dcm-6 hisG4 rfbD1rpsL136 dam13::Tn9 xylA5 mtl-1 thi-1 mcrB1 hsdR2* | New England Biolabs |
| ***H. volcanii* strains** |  |  |
| H26 | DS70 Δ*pyrE2* | (24) |
| H1207 | Δ*pyrE2* *pitA<sub>Nph</sub>* Δ*mrr* | (30) |
| KS4 | H26 Δ*glpK* | (7) |
| KM01 | KS4 *pitA<sub>Nph</sub>* | This study |
| **Plasmids** |  |  |
| pTA131 | Ap<sup>r</sup>; pBluescript II containing *Pfdx:pyrE2* | (24) |
| pJAM202c | Ap<sup>r</sup> Nv<sup>r</sup>; control plasmid derived from pBAP5010 | (64) |
| pTA1106 | Ap<sup>r</sup>; pTA131 with *pitA<sub>Nph</sub>* gene replacement | (30) |
| pJAM2666 | Ap<sup>r</sup> Nv<sup>r</sup>; pJAM2055 with P2<sub>*rrn*</sub>-*glpK-strepII* (expressing GK-StrepII) | (7) |
| pJAM503 | Ap<sup>r</sup> Nv<sup>r</sup>; pBAP5010 with P2<sub>*rrn*</sub>-*His6-panA* | (64) |
| pJAM4351 | Ap<sup>r</sup> Nv<sup>r</sup>; pJAM503-derived with P2<sub>*rrn*</sub>-*his6-glpK* (expressing His-GK) | This study |

<sup>a</sup>Ap<sup>r</sup>, ampicillin resistance; Nv<sup>r</sup>, novobiocin resistance; *larC* encoding LarC (HVO\_2381, UniProt D4GWM9); *glpK* (*hvo\_1541*, UniProt D4GYI5); StrepII tag includes linker as follows: Gly-Thr-Trp-Ser-Pro-Gln-Phe-Glu-Lys; *pitA<sub>Nph</sub>*, *Natronomonas pharaonic* *pitA* (*np\_2262a*, UniProt Q3IRM1) was used to replace *H. volcanii* *pitA* (*hvo\_1871*, UniProt D4GSX4).

### Plasmid construction

To construct the plasmids listed in Table 1, high-fidelity, double-stranded DNA products were generated by polymerase chain reaction (PCR) using Phusion DNA polymerase (New England Biolabs). The primer pairs used for each PCR are listed in Table 2. *H. volcanii* genomic DNA, extracted as described in *The Halohandbook* (23) was used as the PCR template. All PCR reactions followed the manufacturer’s instructions, with 3 % (v/v) DMSO added to the mix. PCRs were performed using an iCycler or MyCycler system (Bio-Rad Laboratories, Hercules, CA). A touchdown-PCR program was utilized to amplify the glycerol kinase gene, *glpK* (*hvo_1541,* UniProt D4GYI5). This program began with an initial denaturation at 98 °C for 2 min, followed by two cycling blocks. The first block comprised 10 cycles of denaturation at 98 °C for 10 sec, annealing at 72 °C for 30 sec (decreasing 0.5 °C per cycle), and extension at 72 °C for 1 min. The second block included 25 cycles of denaturation at 98 °C for 10 sec, annealing at 67 °C for 30 sec, and extension at 72 °C for 1 min. A final extension step at 72 °C for 5 min concluded the reaction. DNA fragments were separated using 0.8 % (w/v) agarose gels containing ethidium bromide (0.5 μg·mL⁻¹) in 1× TAE electrophoresis buffer. Molecular weight standards (GeneRuler 1 kb Plus, Thermo Scientific, Waltham, MA) were used for size estimation. Gels were imaged with a Mini Visionary Imaging System (FOTODYNE, Hartland, WI). Before enzymatic modifications, PCR products were purified using the Monarch PCR and DNA Cleanup Kit (New England Biolabs). Plasmid DNA was isolated from *E. coli* strains using the PureLink Miniprep Kit (Invitrogen, Carlsbad, CA). Following the manufacturer’s protocols, inserts and vector DNA were digested with restriction enzymes (BlpI, NdeI, EcoRI, or XbaI). When necessary, digested plasmids were extracted from agarose gel slices using the Monarch DNA Gel Extraction Kit (New England Biolabs). Digested DNA fragments were ligated with T4 DNA ligase. Inverse PCR products were treated with DpnI and KLD Enzyme Mix per the supplier’s recommendations (New England Biolabs). The fidelity of all cloned PCR-amplified products was verified through Sanger automated DNA sequencing (Eurofins Genomics, Louisville, KY).

**Table 2.**
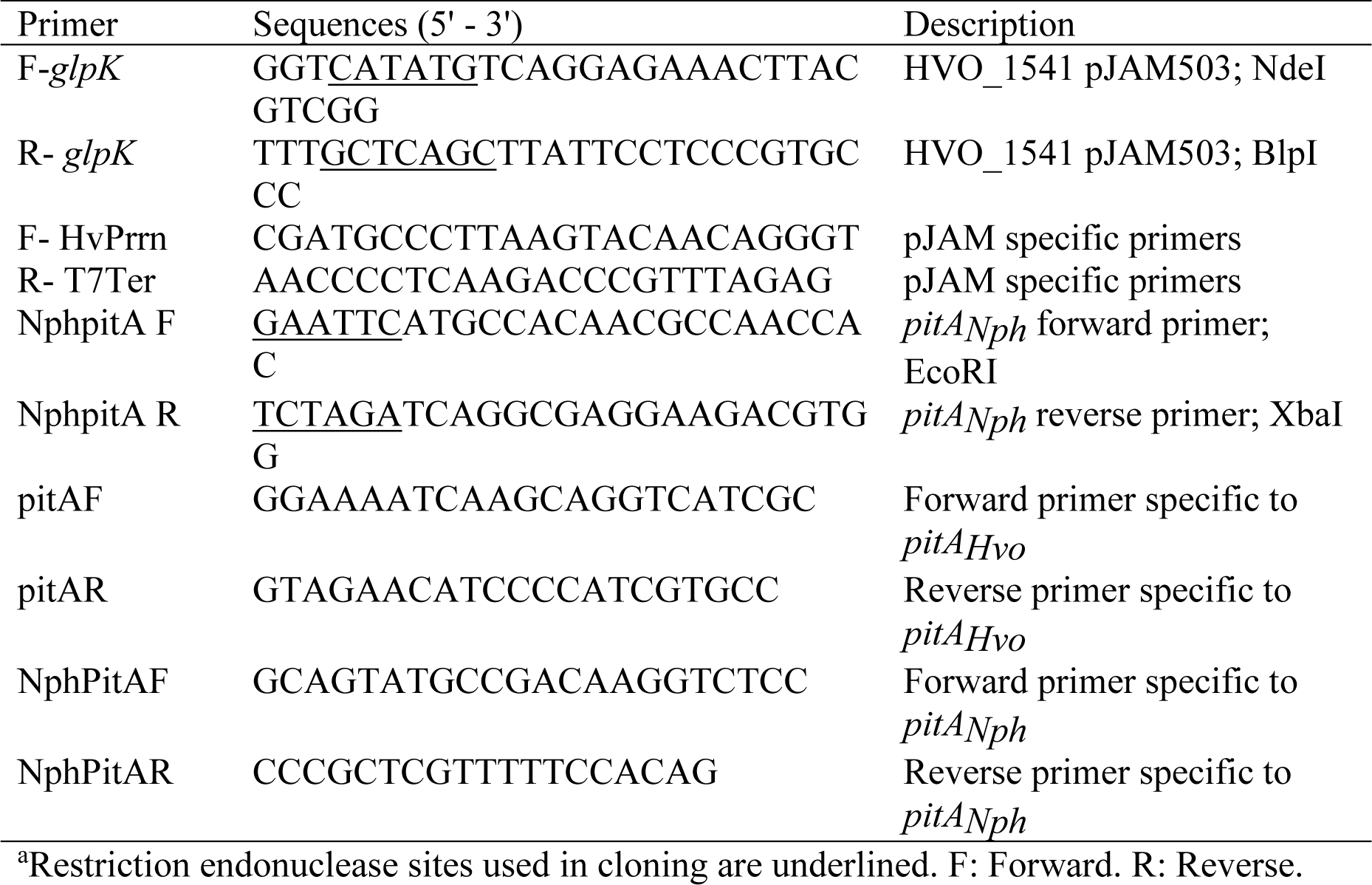
List of primers used in this study^a^.

### Construction of *H. volcanii* strains

To construct *H. volcanii* KM01, the *pitA* (*hvo_1871*, D4GSX4) gene of *H. volcanii* KS04 (H26 *ΔglpK*) was targeted for replacement with the *Natronomonas pharaonis pitA* (*pitA_Nph_*) gene using the *pyrE2*-based “pop-in/pop-out” method (24, 25). In this approach, KS04 was transformed with plasmid pTA1106 prepared in *E. coli* GM2163. *H. volcanii* transformants were plated on Hv-CA agar medium lacking uracil, and growth was counter-selected on media containing 5-FOA. The 5-FOA resistant colonies were screened for the identification of strains with desired replacement of the *H. volcanii pitA* with *pitA_Nph_*. PCR screening of colonies was performed using internal primers (*pitAF* and *pitAR*) specific to the target gene. Colonies that lacked the PCR product were further analyzed using primers (*NphPitAF* and *NphPitAR*), designed to anneal to the flanking regions. The resulting PCR products were resolved by gel electrophoresis and analyzed by DNA sequencing to ensure the accuracy of the KM01 strain.

### Complementation assay

The biological function of affinity tagged GK variants was performed by complementation assay as previously described (7). The GK enzymes fused to an N-terminal polyhistidine tag (His-GK) and C-terminal StrepII tag (GK-StrepII) were compared. In brief, the parent H26, the mutant KS4 (H26 *ΔglpK*), and the mutant carrying either the plasmid expressing His-GK (KS4-pJAM4351), GK-StrepII (KS4-pJAM2666), or the empty vector (KS4- pJAM202c) were cultured on GlyMM and FMM plates (42 °C, 5 days). Growth patterns were compared and imaged using an iBright Imaging System (Invitrogen). The data were interpreted based on the previous finding that *H. volcanii* requires GK when grown on GlyMM compared to FMM (7).

### Protein purification and analysis

*H. volcanii* H1207 strains expressing GK-StrepII (pJAM2666), His-GK (pJAM4351) or empty vector (pJAM202c) were used for protein purification. The *H. volcanii ΔglpK* mutant expressing His-GK (KS04-pJAM4351) was also used for purification. The strains were grown in GlyMM. Cells were grown to stationary phase (1.2 OD_600_, 1 OD_600_ unit equals approximately 10^9^ CFU·mL^-1^) in either 100 mL cultures in 500 mL flasks or 750 mL cultures in 2.8 L Fernbach flasks (200 rpm, 42 °C, 5 days). Cells were harvested by centrifugation (25 min, 5000 × *g*) at 21 °C (room temperature, RT). Cell pellets were stored at -80 °C until use.

For purification of GK-StrepII, cell pellets (from 750 mL cultures) were resuspended in lysis buffer: 20 mM Tris-HCl, pH 7.5, 2 M NaCl, 1 mM CaCl_2_, 3 mM MgCl_2_, 1 mM TCEP, 10 μg/mL DNase I, and EDTA-free protease inhibitor (1 tablet/10 mL). For each gram of wet cell pellet, 5 mL of lysis buffer was used. Cells were disrupted using a French pressure cell with four passes (20,000 lb·in⁻², minimum high ratio of 140, Glen-Mills, NJ, U.S.). The lysate was centrifuged to remove debris (30 min, 13,000 × *g*, 4 °C), and the supernatant was filtered sequentially through 0.45 μm and 0.22 μm pore-size filters. The GK-StrepII protein was purified using Strep-Tactin Superflow Plus Resin (binding capacity 9 mg protein/mL, U.S. Cat. No. 1057978 Qiagen, Germantown, MD). The resin (500 μL resin bed volume) was transferred as a 1 mL slurry to a 5 mL centrifuge column (U.S. Cat. No. 89897 Pierce, ThermoFisher Scientific) and equilibrated with binding buffer: 20 mM Tris-HCl, pH 7.5, 2 M NaCl, 1 mM TCEP. Clarified lysate was applied to the equilibrated resin, and the mixture was incubated at 4 °C for 1 h with gentle rocking. After incubation, the column was centrifuged (500 × *g*, 1 min, RT), and the flow- through was collected. The resin was washed twice with 10 bed volumes of wash buffer: 20 mM Tris-HCl, pH 7.5, 2 M NaCl, 1 mM TCEP. Bound protein was eluted with two-bed volumes of elution buffer: 20 mM Tris-HCl, pH 7.5, 2 M NaCl, 1 mM TCEP, and 5 mM desthiobiotin.

Elution was performed by rotating the resin at 4 °C for 30 min, followed by centrifugation (500 × *g*, 1 min, RT).

For purification of His-GK, cells pellets (from 100 mL cultures) were resuspended in 1 mL of lysis buffer: 20 mM Tris-HCl or 50 mM HEPES, pH 7.5, 2 M NaCl, 5 mM β-mercaptoethanol, 40 mM imidazole, DNase, and EDTA-free protease inhibitor (1 tablet/10 mL). Cells were lysed in 1.8 mL microcentrifuge tubes by sonication on ice (four pulses of 10 s each) (Sonifier Cell Disruptor, Model W185, Heat Systems-Ultrasonics, Inc., Plainview, NY). The lysate was centrifuged (10 min, 13,000 × *g*, 4 °C). The supernatant was applied to Ni-NTA His-Bind resin (100 μL slurry/50uL bed volume, binding capacity 5–10 mg protein/mL; U.S. Cat. No. 70666-3 Millipore Sigma, Burlington, MA) preequilibrated in wash buffer: 20 mM Tris-HCl or 50 mM HEPES, pH 7.5, 2 M NaCl, 40 mM imidazole. The mixture was incubated for 1 h at 4 °C with gentle rocking. The resin was washed three times with 1 mL of wash buffer, and the His-GK protein was eluted three times with 100 μL of elution buffer: 20 mM Tris-HCl or 50 mM HEPES, pH 7.5, 2 M NaCl, 250 mM imidazole.

### Protein quantification

Protein concentrations were measured using the Bradford assay (26) with bovine serum albumin (Bio-Rad Laboratories) as the standard. The assay was performed with the 96-well microplate format with a reaction volume of 250 μL. For each measurement, 5 μL of protein sample was mixed with 250 μL of Bradford reagent, and the mixture was incubated for 5 min at RT. Absorbance at 595 nm (A_595_) was recorded using a microplate reader (EPOCH2, BioTek, Agilent Technologies, Santa Clara, CA). The assay exhibited linearity within the 0 to 2 mg·mL⁻¹ protein range.

### SDS-PAGE

Protein purity and subunit molecular mass were assessed by reducing SDS–PAGE with 12 % polyacrylamide gels, run in Tris-glycine-SDS buffer at 120 V for 2 h. Protein fractions were prepared for electrophoresis by mixing equal volume of Laemmli SDS sample buffer: 100 mM Tris-HCl (pH 6.8), 10 % (v/v) β-mercaptoethanol, 2 % (w/v) SDS, 10 % (v/v) glycerol, and 0.6 mg/mL bromophenol blue. The samples were boiled for 10 min, chilled on ice for 5 min, and centrifuged at 13,000 × *g* for 10 min. Precision Plus Protein Kaleidoscope molecular mass marker (Bio-Rad Laboratories) was used as the standard. The gels were stained with Coomassie Blue and imaged using an iBright Imaging System (Invitrogen) according to the manufacturer’s protocol.

### Immunoblotting analysis

After SDS-PAGE, proteins were transferred to PVDF membranes (0.2 µm) (Amersham, Little Chalfont, UK) at 30 V for 14 h at 4 °C with constant stirring as per standard protocol (Bio-Rad Laboratories). After protein transfer, the membranes were incubated with gentle rocking for 2 h at RT in blocking buffer composed of 5 % (w/v) skim milk powder in TBST: 0.05 M Tris-HCl, pH 7.6, 0.15 M NaCl, 0.1 % (v/v) Tween 20. To detect His-GK, HRP-conjugated 6×His-tag mouse monoclonal antibody (Proteintech Group, Inc, Rosemont, IL, U.S. Cat. No. HRP-660005) was diluted 1:10,000 in blocking buffer and incubated with the membrane for 1 h at RT. Following incubation, the membrane was washed five times with TBST for 5 min per wash. The Amersham ECL Prime substrate mixture (1:1) was applied to the membrane and incubated for 5 min prior to imaging with the iBright Imaging System (Invitrogen) according to the manufacturer’s protocol. To detect GK-StrepII, the rabbit anti-StrepII polyclonal “NWSHPQFEK” antibody (Genescript, Piscataway NJ, U.S. Cat. No. A00626), diluted to 0.25 μg/μL in the blocking buffer, was incubated with the membrane for 1 h at RT. After washing the membrane five times with TBST, the secondary antibody, goat anti-rabbit HRP-conjugated (Southern Biotech, Birmingham, AL, U.S. Cat. No. 4010-05), was diluted 1:10,000 in blocking buffer and incubated with the membrane for 1 h at RT. Following secondary antibody incubation, the washing step was repeated. The Amersham ECL Prime substrate application and iBright Imaging were as described for His-GK.

### Size exclusion chromatography

Prior to size exclusion chromatography (SEC), protein samples were dialyzed using mini D-Tube Dialyzers with a molecular weight cutoff of 12–14 kDa according to supplier (Novagen, Madison, WI). Samples were dialyzed (12–16 h, 4 °C) against buffer (2 L per mL sample) composed of 50 mM HEPES, pH 7.5, 2 M NaCl, and 1 mM DTT with or without 10 % (v/v) glycerol. After dialysis, protein samples (100 μL per run) were applied to a Superdex 200 Increase 10/300 GL (Sigma-Aldrich, St. Louis, MO, U.S. Cat. No. GE28-9909-44), pre-equilibrated in the dialysis buffer. The chromatography was performed in the dialysis buffer at a 0.3 mL·min⁻¹ flow rate. Molecular mass standards included vitamin B12 (1.35 kDa), myoglobin (horse, 17 kDa), ovalbumin (chicken, 44 kDa), γ-globulin (bovine, 158 kDa), and thyroglobulin (bovine, 670 kDa), and the void volume marker Blue Dextran (2,000 kDa) (Bio-Rad Laboratories). Protein elution was monitored by UV absorbance at 280 nm (A₂₈₀) and quantified using the Bradford assay (Bio-Rad Laboratories). Molecular mass estimations were derived from the linear regression (*R*² > 0.99) of the logarithmic values of molecular mass against the gel phase distribution coefficient (K_av_). K_av_ was calculated using the equation: K_av_=(V_R_−V_o_)/(V_c_−V_o_) where V_R_ represents the retention (elution) volume of the protein, V_o_ is the void volume of the column, and V_c_ is the geometric bed volume.

### GK activity assay

GK activity was determined using a coupled assay in which the reduction of NAD^+^ to NADH was monitored by UV-Vis spectroscopy. In brief, GK catalyzes the ATP-dependent conversion of glycerol to glycerol-3-phosphate (G3P). This reaction is quenched and then G3P dehydrogenase (G3PDH) is used to detect the oxidation of G3P to DHAP by monitoring the reduction of the electron acceptor NAD^+^ to NADH at absorbance 340 nm (A_340_). Purified His-GK was used for all activity assays. Reaction mixtures (1 mL) containing GK (0.84 µg protein, equivalent to 0.3 µM homodimer), 3.5 mM MgCl₂·6H₂O, 3.5 mM ATP, 4.6 mM glycerol, 50 mM HEPES, pH 8.0, and 0.1 M NaCl were incubated at 57 °C for 30 min unless otherwise indicated. After incubation, reaction mixtures (200 µL aliquots) were withdrawn and terminated by addition of equal volume of 0.2 N H₃PO₄. The samples were centrifuged at 12,000 × *g* for 10 min at RT to remove precipitated proteins. The second reaction was carried out as previously described (27). The G3P content of 110 µL portions was assayed enzymatically in a total reaction volume of 1 mL. The reaction contained 0.011 N NaOH for neutralization, followed by 1.1 mM NAD^+^, 0.66 M hydrazine sulfate (adjusted to pH 9.4 with NaOH), 1 % (w/v) nicotinamide-sodium carbonate buffer, and rabbit muscle G3P dehydrogenase (8-10 U; Sigma, EC 1.1.1.8, 500 U). The reaction was incubated for 1 h at 30 °C. NADH formation was measured at A_340_. Product formation was quantified using G3P standards ranging from 0 to 5 mM, demonstrating a linear relationship in assays (without GK), as confirmed by linear regression analysis (R² > 0.99).

### GK activity optimization

Influence of pH, temperature, salinity, and ions (monovalent and divalent) on GK activity were tested by varying one parameter at a time while keeping others constant. The effect of magnesium concentration was tested across a range of 0 to 500 mM (0, 0.5, 3.5, 10, 20, 50, 100, and 500 mM MgCl_2_). Additionally, the effect of different cations on GK activity was evaluated by comparing the influence of KCl, LiCl, CaCl₂, MnCl₂, ZnSO_4_, CoCl_2_, and MgCl₂ at a concentration of 3.5 mM. The effect of buffer and pH was tested using MES (pH 6.0, 6.5), HEPES (pH 7.0, 7.5, 8.0, 8.2), CAPS (pH 9.0, 9.5, 10.0), TES (pH 7.0, 7.5, 8.0, and 8.2), and Tris-HCl (pH 7.0, 7.5, 8.0, and 8.2) buffers. Temperature was tested from 7 °C to 100 °C (in 5 °C increments up to 97 °C, with a final test at 100 °C). Salinity was tested from 0.1 to 4 M NaCl, with 0.1 M, 0.15 M, and 0.5 to 4 M NaCl concentrations in 0.5 M intervals. The salinity from the first reaction was diluted to a concentration that had no influence on the commercial G3PDH used in the second reaction (as confirmed experimentally). The effect of organic solvent (DMSO) was also tested at concentrations of 5 % and 10 % (v/v) DMSO with NaCl at 0.1, 1, 2, 3, and 4 M. For these experiments, the salinity carryover of the G3P product was equalized to 0.44 M in the second reaction, as this corresponded to the highest salinity condition tested.

### GK kinetic analysis and testing of fructose-1,6-bisphosphate as an allosteric effector

Kinetic analysis of GK was performed under the optimal conditions of pH (8.0), temperature (57 °C), and salinity (100 mM NaCl). The concentration of each substrate and magnesium cofactor was varied to assess their impact on enzyme activity: magnesium (0, 0.05, 0.1, 0.15, 0.2, 0.3, 0.4, 0.5, 1.0, 2.0, and 3.5 mM MgCl_2_), glycerol (0, 0.2, 0.4, 0.6, 0.8, 1, 2, 3, and 4.6 mM), and ATP (0, 0.2, 0.4, 0.6, 0.8, 1, 2, and 3.5 mM). Kinetic parameters were compared using the Michaelis-Menten equation, Lineweaver-Burk plot, and Hill equation to analyze the enzyme’s behavior for each substrate and magnesium cofactor. Kinetic analysis was based on three experimental replicates with three technical replicates each. The potential inhibitory effect of fructose-1,6- bisphosphate (FBP), a known allosteric inhibitor of bacterial glycerol kinase, was evaluated using the *H. volcanii* GK. Enzymatic activity assays were performed in the presence and absence of 20 mM FBP in the reaction mixture to assess any changes in activity levels.

### Degradation of crude glycerol by His-GK

To evaluate the ability of *H. volcanii* GK to degrade crude glycerol, enzymatic activity assays were performed using crude glycerin as the substrate. The crude glycerin sample contained 87 % glycerol, 4.2 % water, 4.5 % ash, and 1.3 % methanol. The amount of crude glycerin used in the assay corresponded to the glycerol concentration typically utilized in activity assays, at a final concentration of 4.6 mM glycerol.

The enzymatic activity with crude glycerin was measured and compared to activity with pure glycerol under identical assay conditions to determine the relative degradation efficiency.

### GK tolerance to freeze-thaw, salinity and temperatures

To examine the influence of freeze-thaw, the purified His-GK was dialyzed against 50 mM HEPES, pH 7.5, 2 M NaCl, and 1 mM DTT buffer containing 10 % (v/v) glycerol. Enzymatic activity was measured on the freshly prepared protein. The sample was then stored at -80 °C for 5 days. After storage, the sample was thawed, and the enzymatic activity assay was repeated to compare the GK activity before and after freezing. The stability of GK in different saline conditions was assessed by resuspending His-GK to a final concentration of 17 µg/mL in buffer containing 50 mM HEPES (pH 8) and NaCl at final concentrations of 0.1, 0.15, 1, 2, 3, and 4 M. The enzyme samples in the respective buffers were stored at 4°C, and enzymatic activity was measured at 0, 24, 48, and 72 h. For these enzymatic activity assays, the final GK concentration in each reaction was 0.84 µg/mL (equivalent to 0.3 µM homodimer). To isolate the effects of storage stability rather than salinity on enzymatic activity, the NaCl concentration in the first reaction of all conditions was standardized to 0.2 M by adding NaCl as needed. Temperature stability was tested under two salinity conditions: 0.1 M NaCl and 2 M NaCl. His-GK was diluted to 17 µg/mL in 50 mM HEPES (pH 8) with the respective salinity concentration and incubated at 24°C, 33°C, 42°C, 51°C, 57°C, 60°C, and 65°C for 1 h. After incubation, the enzymatic activity assay was performed under optimal conditions at 57°C. The final NaCl concentrations in the enzymatic reactions were 5 mM for the 0.1 M NaCl buffer and 100 mM for the 2 M NaCl buffer. A no-enzyme reaction was included as a blank for all measurements.

### Differential scanning fluorimetry (DSF) thermal shift assay

His-GK was purified from KM01-pJAM4351 grown in GlyMM and stored at -80°C in a buffer containing 10 % (v/v) glycerol. For the thermal shift assay, the enzyme was diluted in a buffer containing 50 mM HEPES, pH 7.5, 2 M NaCl, and glycerol at 1 % or 10 % (v/v) final concentration, as indicated. Reactions were prepared in a total volume of 25 µL. The final enzyme concentration was 2.9 µM, and Sypro Orange Protein Gel Stain (5000× in DMSO, Invitrogen U.S. Cat. No. S6650) was added to a final concentration of 2.5×. Samples were blanked against a non-enzyme control containing the respective glycerol concentration to account for background fluorescence. The thermal shift assay was conducted using a C1000 thermal cycler with the CX96 real-time system (Bio-Rad Laboratories). Samples were subjected to a temperature gradient from 20°C to 95°C at a rate of 1°C per min. Fluorescence data were recorded continuously to monitor the unfolding process. The fluorescence intensity data were analyzed to identify the melting peak, which represents the temperature at which the protein unfolds. Results were compared between conditions with 1% and 10% glycerol to assess the effect of glycerol concentration on protein stability.

### Protein modeling

The 3D structural models of *H. volcanii* GK (UniProt ID: D4GYI5, HVO_1541) were generated using AlphaFold (28). High-confidence monomer and homodimer models were produced, with intrinsic predicted Template Modeling scores (ipTM) and predicted Template Modeling scores (pTM) of 0.94 and 0.92 for the monomer and 0.83 and 0.85 for the homodimer, respectively. To map the active site, glycerol was overlaid onto the AlphaFold-predicted models using structural alignment with the X-ray crystal structure of substrate-bound GK from *Thermococcus kodakarensis* (PDB ID: 6K79) as a reference. Ligands, including Mg^2+^, ATP, and glycerol, were positioned in the active site, and interactions were visualized using ball-and-stick diagrams. The AlphaFold-predicted Local Distance Difference Test (pLDDT) scores were used to assess the confidence of the monomer model, with a color gradient indicating per-residue reliability. The homodimer model was visualized with Coulombic surface coloring to illustrate electrostatic properties. Visualization and analysis were performed using ChimeraX (29). The N-terminal and C-terminal residues were annotated, and putative active site residues were identified and labeled based on UniProt annotations and spatial proximity to the ligands.

### Statistical Analysis

Enzymatic activity assays were performed under defined conditions (as outlined in earlier section). Each experiment included technical replicates performed in at least triplicate and was conducted a minimum of three times to ensure reproducibility. Data are presented as mean values ± standard deviation (SD). Statistical significance between experimental groups was assessed using Student’s t-test, with a threshold for significance set at p-value < 0.05. All statistical analyses were performed using Microsoft Excel.

## RESULTS

### 3D-structural modeling of H. volcanii GK

AlphaFold modeling was used to predict the 3D- structure of the *H. volcanii* GK and determine the optimal location to attach an affinity tag to facilitate protein purification. The 3D-modeling suggested the *H. volcanii* GK forms a homodimer and enabled prediction of the enzyme active site in this structure including the binding sites for Mg^2+^, ATP and glycerol (**Figure 1**). The C-terminal region appeared near the active site and homodimer interface. By contrast, the N-terminal region was substantially distant from these locations. These results suggested that adding an affinity tag to the C-terminus may cause misfolding or obstruct the active site during purification, whereas tagging the N-terminal region was predicted to have minimal impact on the enzyme.

**Figure 1.**
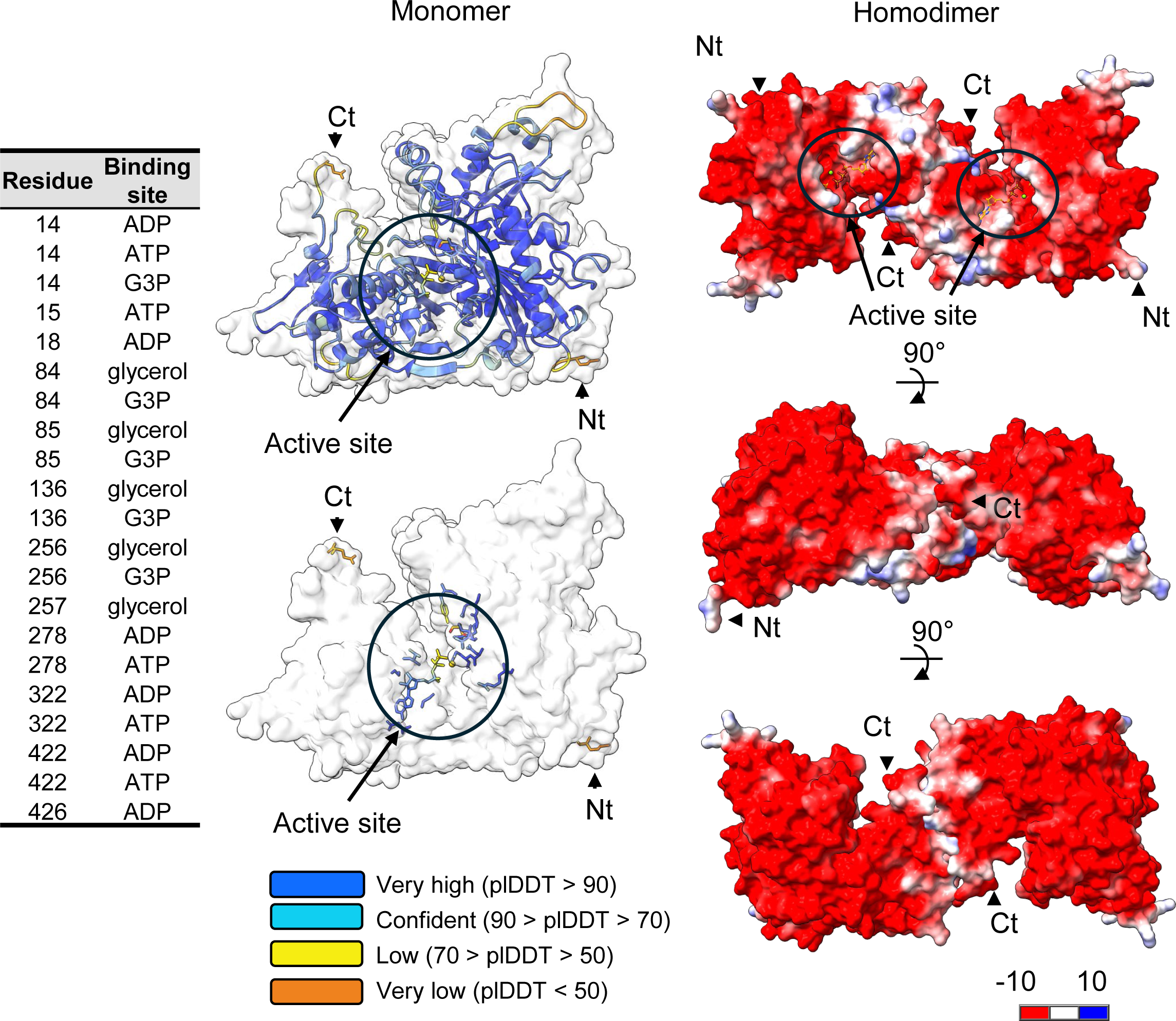
3D structural models of monomeric and dimeric forms of *H. volcanii* glycerol kinase with putative active site residues indicated. *H. volcanii* glycerol kinase (GK, HVO_1541 UniProt D4GYI5) bound to Mg^2+^ and ATP as a monomer (ipTM = 0.94 pTM = 0.92) and homodimer (ipTM = 0.83 pTM = 0.85) was modeled at high confidence using AlphaFold 3 server. Other oligomeric configurations were predicted only at low confidence (ipTM < 0.8). Glycerol was overlayed onto the 3D-models using the X-ray crystal structure of substrate bound GK of *Thermococcus kodakarensis* (PDB:6K79) as a guide. Active site ligands are represented in ball and stick diagram. N-terminal (Nt) and C-terminal (Ct) residues are indicated. Coloring scale of monomer (pLDDT a per-atom confidence) and homodimer (Coulombic surface coloring) indicated. Right list, residue number and putative binding function based on UniProt annotation.

### Biological activity of affinity tagged *H. volcanii* GK

To further analyze the affinity tagging strategy, a complementation assay using a *ΔglpK* (GK) mutant strain was used. The *ΔglpK* (GK) mutant is unable to grow on glycerol unless the *glpK* gene is reintroduced to restore its activity or the cells are provided with an alternative carbon source like fructose (7). Our previous work demonstrated that GK fused to a C-terminal StrepII tag (GK-StrepII) retained *in vivo* functionality using this approach (7). Here we examined the *ΔglpK* mutant expressing the GK fused to an N-terminal His tag (His-GK) from a plasmid. Through the expression of the His-GK, this strain was found to be restored for growth on glycerol to levels comparable to the parent (H26) **(Figure 2A).** By contrast, the *ΔglpK* mutant carrying the empty vector control was unable to grow on glycerol. As an additional control, all strains were shown to be able to grow on fructose (**Figure 2B**). These results are consistent with the essential role of the GK in the metabolism of glycerol compared to other carbon sources and reveal that the N-terminal His-tag did not negatively impact the biological function of the *H. volcanii* GK.

**Figure 2.**
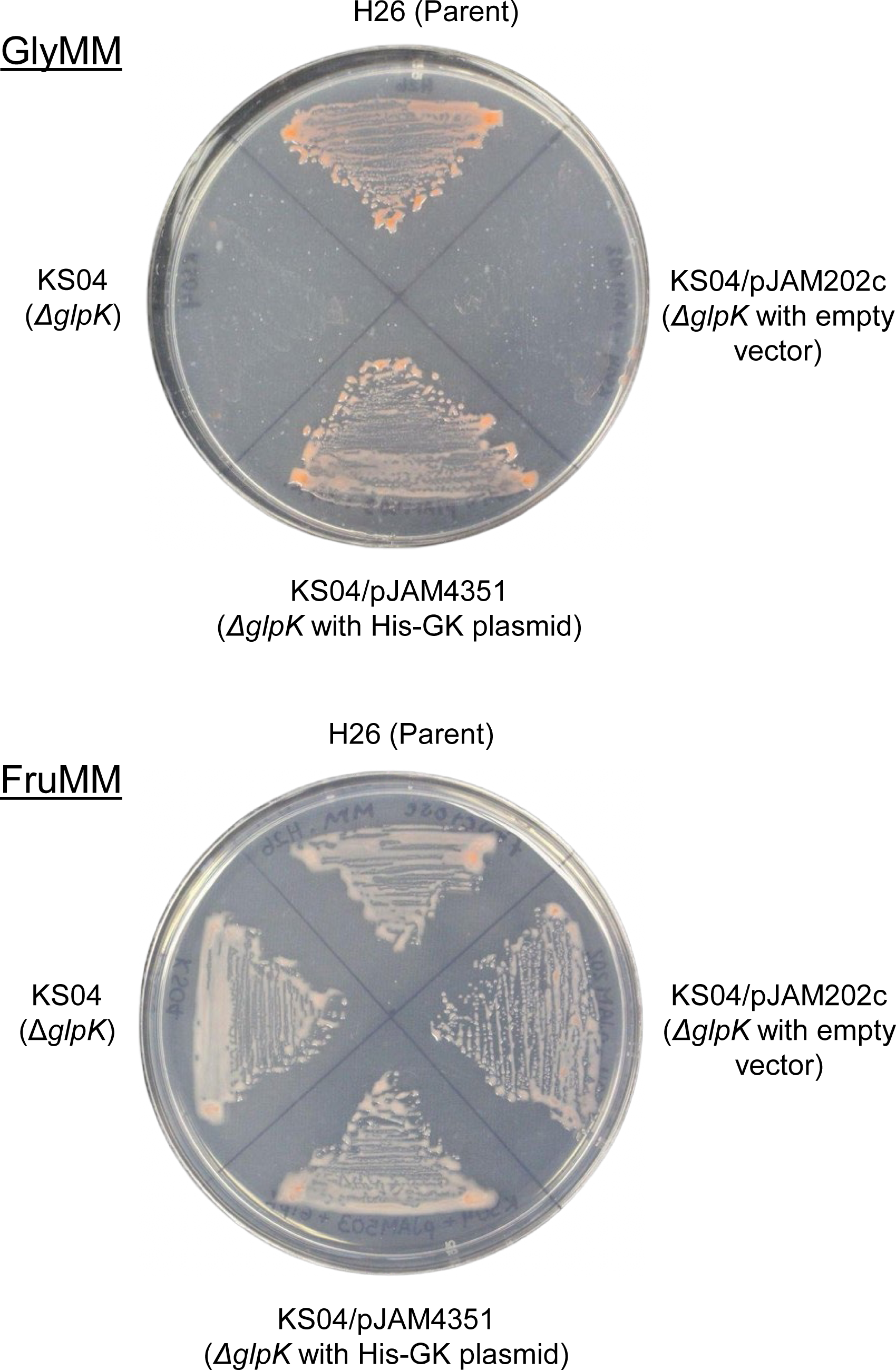
N-terminal His tagged glycerol kinase is biologically active. *H. volcanii* strains were compared for growth on glycerol (GlyMM) and fructose (FMM) as the sole carbon source as indicated. *H. volcanii* strains tested included: H26, parent; KS04, *ΔglpK* mutant; KS04/pJAM202c, *ΔglpK* mutant with empty vector; KS04/pJAM4351, *ΔglpK* mutant with His- GK expression plasmid. His-GK, N-terminal His tagged glycerol kinase.

### N-terminal His-tag is optimal for *H. volcanii* glycerol kinase purification

To determine whether the N-terminal His-tag was optimal for GK purification, *H. volcanii* strain H1207 was used as host to express the His-GK from a plasmid (pJAM4351). The H1207 strain is genetically modified to avoid the non-specific binding of proteins to the Ni^2+^-affinity resin (30). After growth of this expression strain in ATCC974 rich medium, the His-GK protein was purified by Ni^2+^-affinity resin. The GK-StrepII was similarly expressed in H1207 and purified by StrepTactin resin for comparison. By this approach, the His-GK was found to be of 4.5-fold higher purification yield (3 mg of protein per liter of culture or mg/L) than observed for GK- StrepII at 0.66 mg/L. Further analysis revealed His-GK was purified to apparent homogeneity, as determined by SDS-PAGE followed by Coomassie blue staining and immunoblot analysis with anti-His antibodies (**Figure 3A**). These results suggested that use of the N-terminal His-tag would be ideal for *H. volcanii* GK biochemical studies.

**Figure 3.**
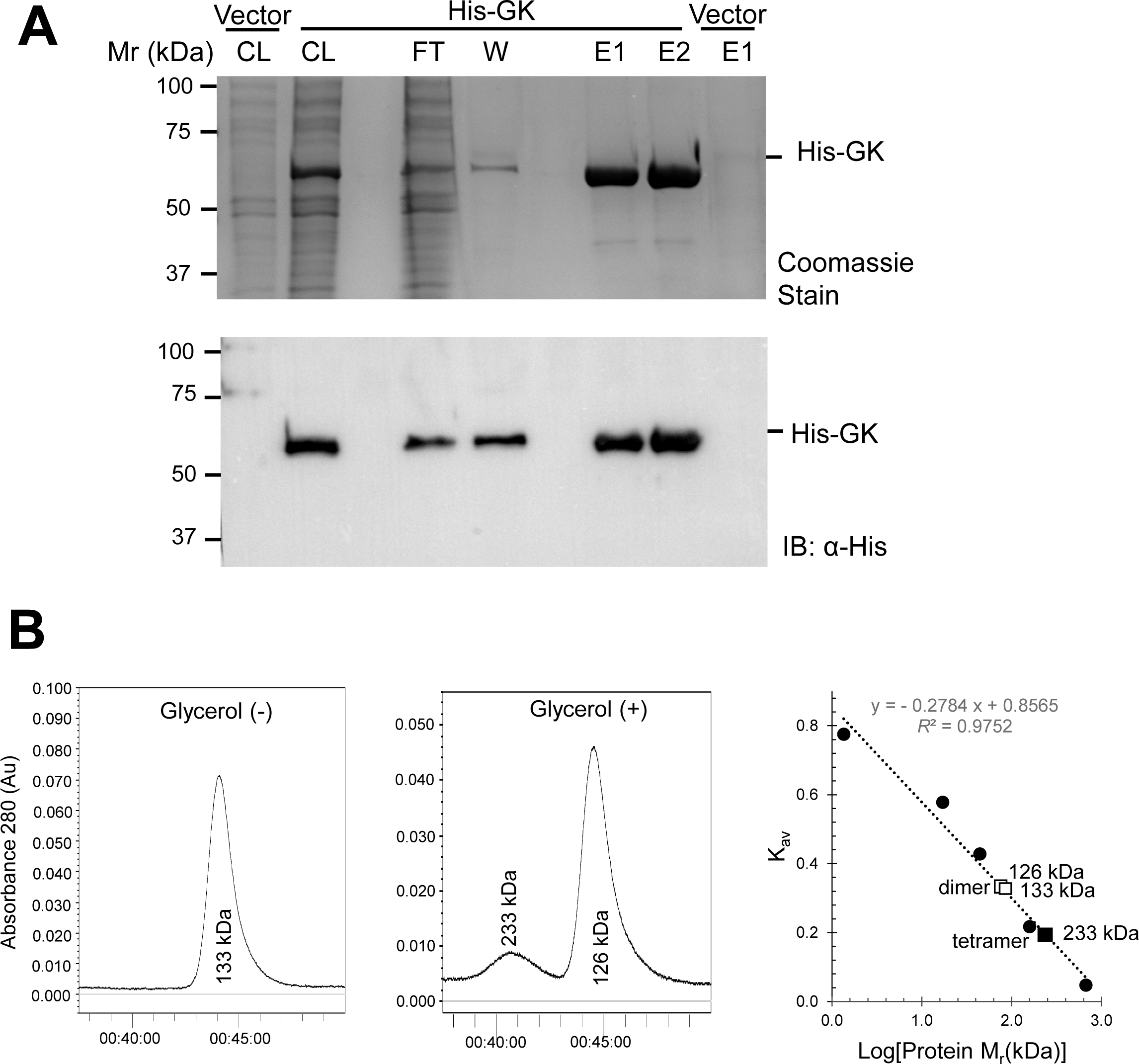
Purification of *H. volcanii* glycerol kinase (His-GK) and glycerol-induced shift from homodimer to homotetramer configuration. A. His-GK is purified to homogeneity as demonstrated by SDS-PAGE. His-GK was synthesized in ATCC974-grown *H. volcanii* H1207-pJAM4351 and purified by Ni^2+^- NTA resin in 50 mM HEPES, pH 7.5, 2 M NaCl buffer. Vector, H1207-pJAM202c with the empty vector control similarly analyzed for comparison. Protein fractions were separated by reducing 12% SDS-PAGE and analyzed by Coomassie stain and immunoblotting using anti-His antibodies (IB: α-His) as indicated. Cell lysate (CL), flow- through (FT), wash fraction (W), and elution fractions (E1 and E2, 2 µg per lane). Left: M_r_, molecular weight standards. B. Glycerol-induced shift of His-GK from homodimer to homotetramer. His-GK was synthesized in GlyMM-grown *H. volcanii* H1207-pJAM4351 and purified by Ni^2+^-NTA resin followed by size exclusion chromatography (SEC) using a Superdex 200 Increase 10/300 GL column. Purification buffer was 50 mM HEPES pH 7.5, 2 M NaCl, 1 mM DTT supplemented with (+) or without (-) 10% (v/v) glycerol as indicated. Chromatographs (left and middle panel) and comparison of molecular mass (M_r_) standards (closed circles) to the His-GK homodimer and homotetramer (open and closed square, respectively) (right panel). M_r_ (*K*_av_) values observed for His-GK at 126 kDa (0.272), 133 kDa (0.265), and 233 kDa (0.197). Calculated theoretical M_r_ values for His- GK monomer (56.7 kDa), homodimer (113.4 kDa), and tetramer (226.8 kDa) for comparison.

### Oligomeric state of *H. volcanii* glycerol kinase affected by glycerol

To further characterize the biochemical properties of the enzyme, His-GK was purified from *H. volcanii* H1207- pJAM4351 grown on glycerol as the sole carbon source (GlyMM). Purification was done using Ni²⁺-affinity resin, followed by SEC to examine the structural conformation of the enzyme. In the absence of glycerol in the purification buffers, the SEC chromatograph showed a single peak corresponding to a GK homodimer (**Figure 3B**). However, when 10% (v/v) glycerol was added to the buffers, two protein peaks were detected, with one corresponding to the homodimer and another to the homotetramer **(Figure 3B)**. These results revealed His-GK to exhibit a glycerol-dependent conformational shift from a dimer to dimer-tetramer equilibrium state.

### Influence of metal ion, buffer, temperature, pH, and salinity on *H. volcanii* glycerol kinase enzymatic activity

To further investigate the biochemical properties and optimize the GK for biocatalysis, its enzymatic activity was assessed using a two-step reaction series. In the first step, glycerol and ATP were converted to G3P by GK. After quenching the reaction, the G3P product was diluted and quantified using G3P dehydrogenase which oxidizes G3P to DHAP using NAD^+^-as the electron acceptor which can be monitored spectrophotometrically. To evaluate the influence of reaction parameters on His-GK activity, one factor was varied at a time, while ensuring the second reaction in the series was not influenced by these alterations. As shown in **Figure 4A**, His-GK required addition of a divalent cation, such as Mg^2+^, for catalytic activity and demonstrated high activity across a wide MgCl_2_ concentration range (10 to 100 mM) with optimal activity at 3.5 mM MgCl_2_. Mn^2+^ and Co^2+^ could substitute for Mg^2+^ in this reaction (**Figure 4B**). By contrast, the divalent cation Zn^2+^ only supported 30% of the activity, while the divalent cation Ca^2+^ and the monovalent cations K^+^ and Li^+^ were unable to serve as replacements (**Figure 4B**). The activity of His-GK was relatively consistent across different buffer types (MES, TES, Tris, HEPES, and CAPSO) and a broad pH range (6.0 to 10.0) **(Figure 4C**). By contrast, a clear temperature preference was observed, with optimal activity occurring at 50-60 °C (**Figure 4D**). Salinity also impacted activity with the His-GK more active at low concentrations (100 to 200 mM) of NaCl, yet still displaying ∼50% activity at 4 M NaCl (**Figure 4E**). With these variables optimized, His-GK was found to display an activity of 143 ± 17.0 µmol G3P/min/mg when assayed in 50 mM HEPES, pH 8.0, 100 mM NaCl buffer supplemented with 3.5 mM MgCl_2_ at 57 °C.

**Figure 4.**
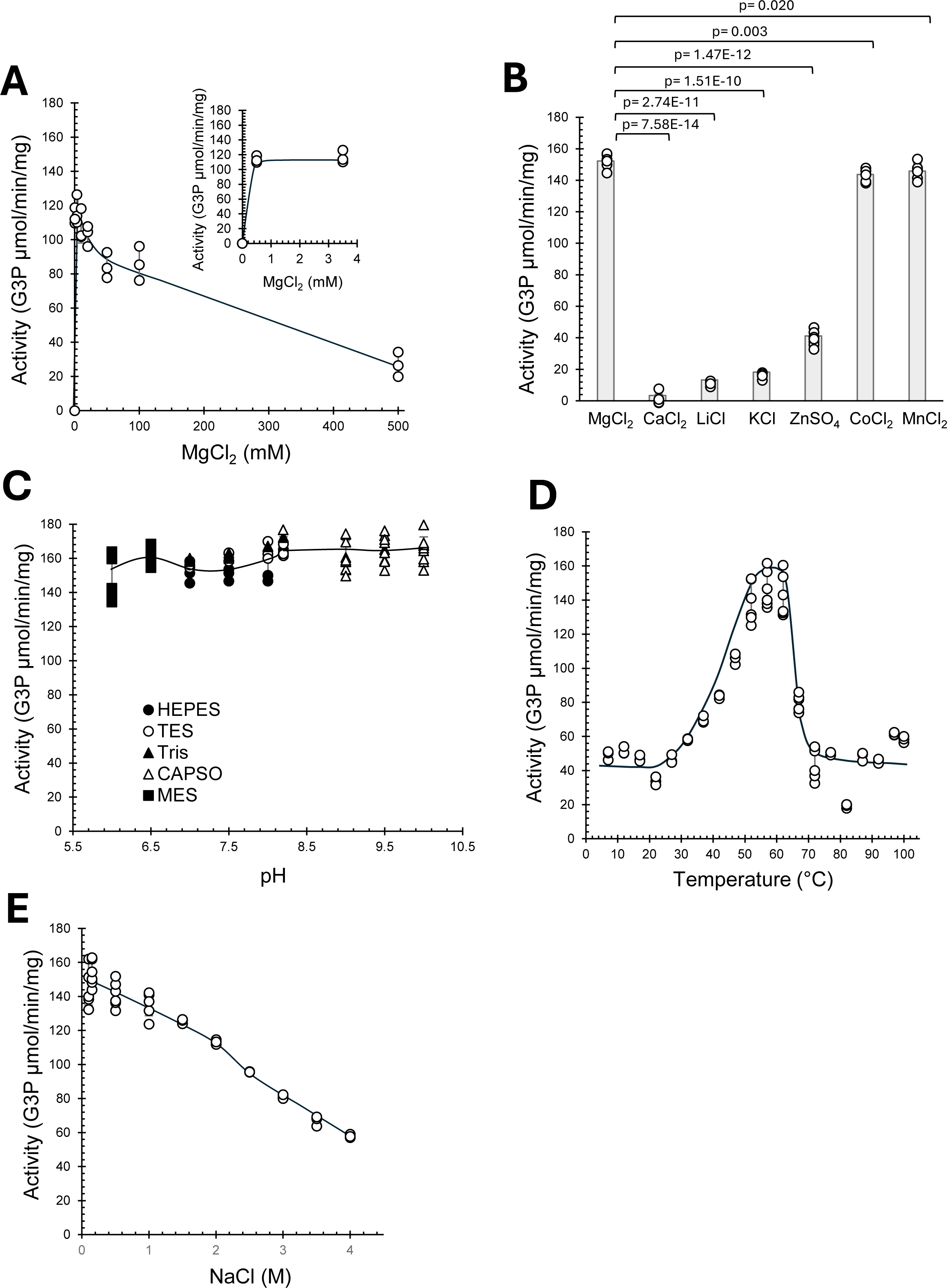
Optimal divalent cation, buffer, temperature and NaCl conditions for *H. volcanii* glycerol kinase enzymatic activity. His-GK enzyme purified from *H. volcanii* KM01-pJAM4351 grown on GlyMM was used for activity assays. A. His-GK activity in concentrations from 0 to 500 mM of MgCl_2_. B. His-GK divalent metal ion requirement. Assay included 3.5 mM MgCl_2_ or replacement of this cationic salt with CaCl_2_ LiCl, KCl, ZnSO_4_, CoCl_2_ or MnCl_2_ as indicated. C. His-GK activity from pH 6.0 – 10.0. Activity was analyzed with HEPES, TES, Tris, MES and CAPSO buffers as indicated. D. His-GK activity at various temperatures. Activity was monitored in intervals of 5°C from 7°C to 97°C and at 100°C, at pH 8.0. E. His-GK activity at different concentrations of NaCl. Activity was measured in 50 mM HEPES, pH 8.0, at 57 °C with NaCl concentrations varied from 0.1 to 4 M NaCl. Statistical significance was evaluated using a t-test, with p < 0.05 considered significant.

### Kinetic parameters and positive cooperativity of *H. volcanii* glycerol kinase

Kinetic analysis revealed His-GK did not follow Michaelis-Menten behavior but instead exhibited positive cooperativity for ATP, glycerol, and magnesium, as indicated by Hill equation parameters **(Figure 5).** The Hill coefficients (n) for ATP (n = 2.35 ± 0.08) and glycerol (n = 2.12 ± 0.19) were consistent with a strong positive cooperativity for the enzyme binding these substrates. The Hill coefficient for Mg^2+^ (n = 1.59 ± 0.15) suggested the cooperativity for this divalent metal cofactor was less pronounced compared to ATP and glycerol. Based on the sigmoidal behavior of the enzyme, dissociation constants (*K*_d_) were determined and found to be 1.6 ± 0.08 mM for ATP, 1.44 ± 0.16 mM for glycerol and 0.089 ± 0.006 mM for Mg^2+^, suggesting the enzyme had a much higher affinity for the Mg^2+^ cofactor when compared to the glycerol and ATP substrates.

**Figure 5.**
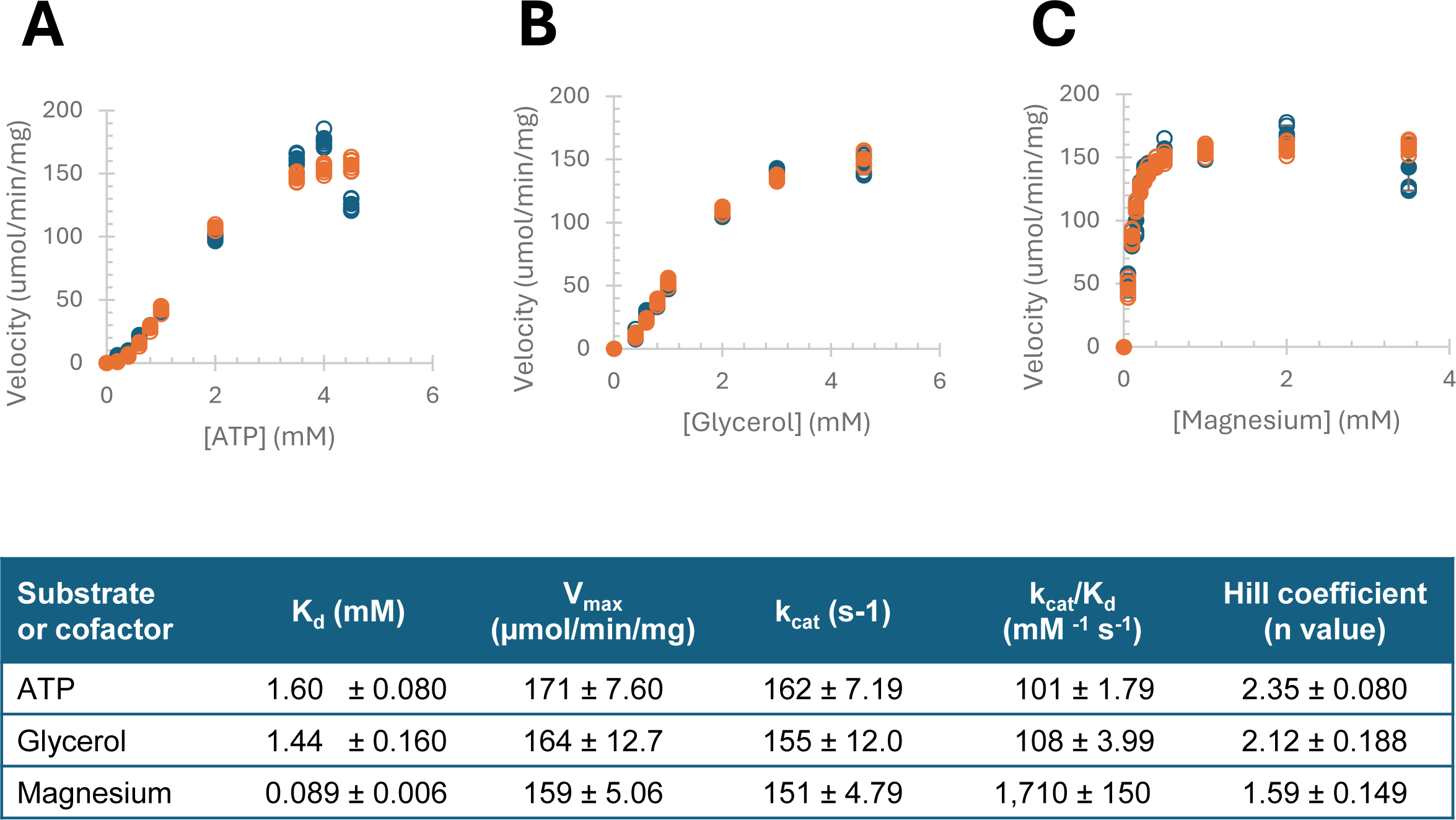
Kinetic analysis of His-GK with the substrates ATP and glycerol, and the cofactor magnesium. His-GK purified from KM01-pJAM4351 grown in GlyMM medium, was used for all kinetic analysis. The activity assay was performed under optimal conditions using 50 mM HEPES buffer, pH 8, 0.1 M NaCl, and 57°C. The enzymatic kinetics show a non-Michaelis Menten behavior, and data were analyzed with the Hill equation. Experimental velocity (blue) and calculated velocity (orange) are presented with ATP (A), glycerol (B) and magnesium (C) as indicated. GK activity was determined using a coupled spectrophotometric reaction with NADH production monitored by absorbance 340 nm (A_340_). Kinetic analysis was calculated based on three experimental replicates with three technical replicates each. The Hill equation used was Y_calculated_ = (Vmax)[L]^*n* / (*K_d_* + [L]^*n*), where Y represents the fraction of occupied binding sites, [L] is the ligand concentration, *n* is the Hill coefficient indicating cooperativity, and *K_d_* is the dissociation constant. Squared residuals (Y_experimental_ − Y_calculated_)^2^ were summed to compute the sum of square residuals (SSR). Excel Solver was employed to minimize SSR and optimize the parameters for the best fit of the data.

The maximum velocity (*V*_max_) and turnover number (*k*_cat_) values were comparable for all three reagents tested averaging 165 ± 8.45 µmol/min/mg and 156 ± 7.99 s⁻¹, respectively. Thus, when the *K*_d_ values were considered to calculate the catalytic efficiency (*k*_cat_/*K*_d_), the substrates ATP and glycerol were found to be 16- to 17-fold lower at 101 ± 1.79 to 108 ± 3.99 mM⁻¹ s⁻¹, when compared to Mg^2+^ at 1,710 ± 150 mM⁻¹ s⁻¹. These results reveal *H. volcanii* GK displays positive cooperativity with its substrates (glycerol and ATP) and has high catalytic efficiency with the magnesium cofactor offering important perspectives on its enzymatic characteristics.

### Effect of organic solvent (DMSO), FBP, and crude glycerol on *H. volcanii* GK activity

The influence of organic solvents was assessed by measuring the catalytic activity of His-GK in the presence of 5 % and 10 % (v/v) DMSO at NaCl concentrations of 0.1, 1, 2, 3, and 4 M. As shown in **Figure 6A**, the addition of 5 % DMSO caused a gradual decline in enzyme activity as salinity increased. In 5 % DMSO, the enzyme retained nearly full activity at 0.1 M NaCl but decreased to approximately half its activity at 4 M salinity. Although His-GK activity decreased at all salinities tested in the presence of 10 % DMSO, the enzyme remained relatively active, maintaining an average of 63 % activity regardless of salinity. To evaluate the potential of FBP as an allosteric inhibitor, His-GK activity was measured in the presence of 20 mM FBP, the highest concentration reported to inhibit the *E. coli* GK (31). No significant inhibition was observed, as His-GK activity levels were comparable to the control reaction without FBP **(Figure 6B).** Additionally, the ability of His-GK to degrade crude glycerin, which contains the organic solvent methanol, was also tested. His-GK demonstrated the same efficiency in degrading crude glycerin as it did with pure glycerol **(Figure 6B)**.

**Figure 6.**
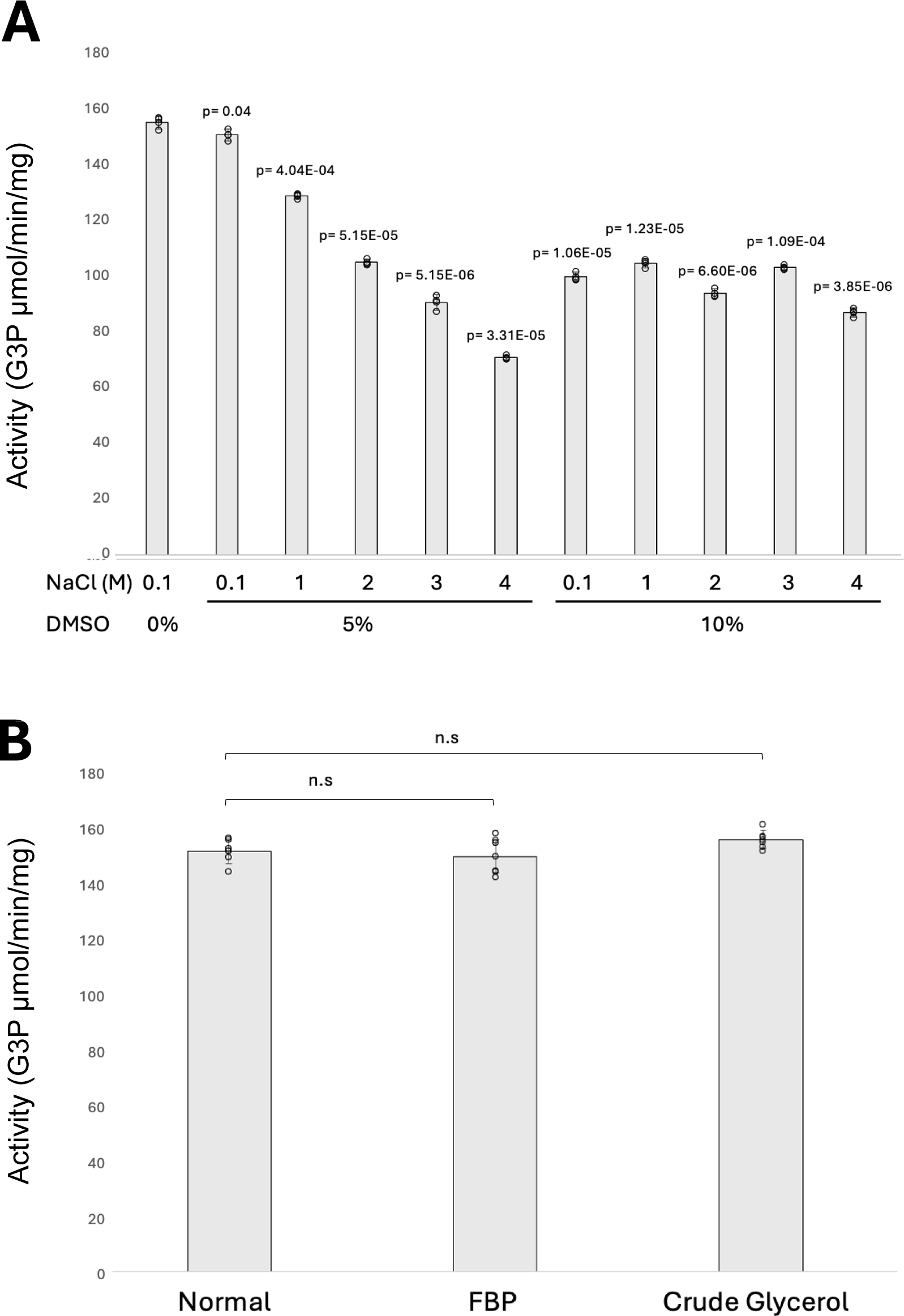
Effect of organic solvent (DMSO), FBP, and crude glycerol on *H. volcanii* GK activity. The His-GK enzyme, purified from *H. volcanii* KM01-pJAM4351 grown on GlyMM, was used for activity assays. A. His-GK activity was evaluated in the presence of 5% and 10% (v/v) DMSO at NaCl concentrations of 0.1, 1, 2, 3, and 4 M. B. The potential inhibitory effect of FBP was tested by adding 20 mM FBP directly to the reaction mixture. Enzymatic degradation of crude glycerol was performed using crude glycerin as the substrate. The enzymatic activity with FBP and crude glycerin was measured and compared to a control reaction where pure glycerol served as the substrate without FBP, under identical assay conditions, to determine the relative degradation efficiency. Statistical significance was evaluated using a t-test, with p < 0.05 considered significant and p > 0.05 denoted as not significant (n.s.).

### Stability of His-GK activity at varying temperatures and salinity levels

To evaluate the stability of His-GK to freeze-thaw, the protein was stored in 50 mM HEPES, pH 7.5, 2M NaCl buffer containing 10 % (v/v) glycerol at -80 °C for five days. After thawing, enzymatic activity was reassessed. Remarkably, 100 % of the enzymatic activity was retained after storage at -80 °C, with slightly higher activity observed compared to the protein stored at 4 °C. These findings indicate that storage at -80 °C in glycerol supplemented buffers effectively preserves His-GK activity, ensuring protein function and stability over time. To assess thermal stability, His-GK was incubated for 1 h at 24 °C, 33 °C, 42 °C, 51 °C, 57 °C, 60 °C, and 65 °C in 50 mM HEPES buffer, pH 8, containing either 2 M NaCl **(Figure 7A)** or 100 mM NaCl **(Figure 7B)**. Enzymatic activity was determined with the optimal parameters of 50 mM HEPES buffer, pH 8, and 57 °C, the final reaction NaCl concentrations were of 100 or 5 mM, respectively. Remarkably, His-GK retained 100 % of its enzymatic activity across all tested temperatures when incubated in the 2 M NaCl supplemented buffer **(Figure 7A).** In contrast, under low salinity (100 mM NaCl), the enzyme remained stable up to 51 °C but showed reduced activity at higher temperatures, retaining only 36 % activity at 57°C. At 60 °C and 65 °C, the enzyme was fully inactivated **(Figure 7B).** These findings suggest that long-term incubation at elevated temperatures requires high-salinity buffers to effectively preserve His-GK activity and ensure its stability over time. To evaluate His-GK stability under varying salinity conditions, enzymatic activity was measured after incubation in 50 mM HEPES buffer, pH 8, with NaCl concentrations of 0.1, 0.15, 1, 2, 3, and 4 M. Samples were stored at 4 °C, and enzymatic activity was assessed at 0, 24, 48, and 72 h. Activity was reassessed with the optimal parameters of 50 mM HEPES buffer, pH 8, and 57°C, with a final reaction NaCl concentration of 0.2 M, ensuring all salinities were equalized during the enzymatic activity assay. The results demonstrated excellent stability across all tested salinities, with most conditions retaining 100 % of enzymatic activity over time. The exception was 4 M NaCl, which showed a slight decrease in activity but still retained more than 80 % activity after 72 h **(Figure 7C)**.

**Figure 7.**
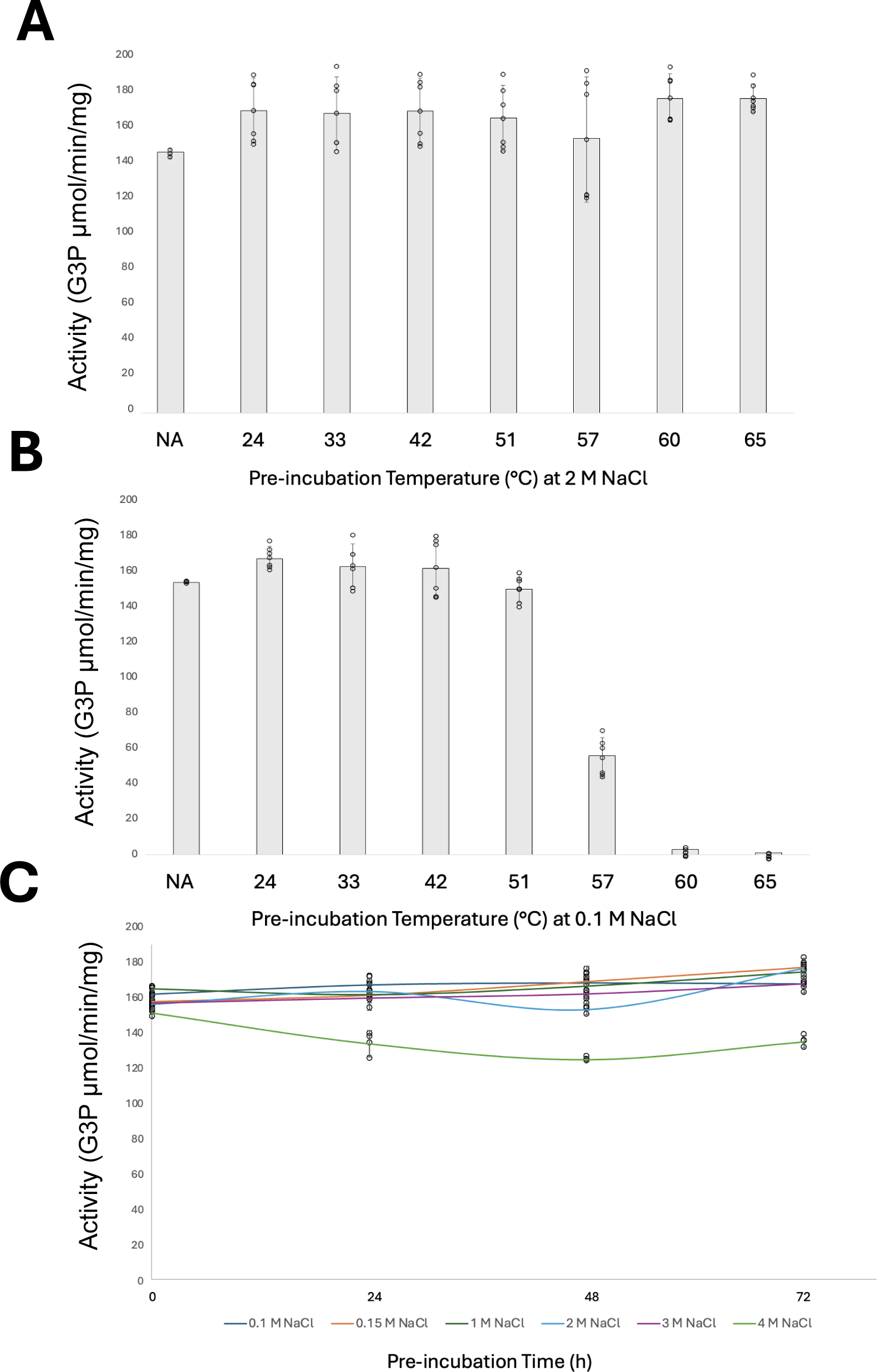
Stability of His-GK activity at varying temperatures and salinity levels. His-GK enzyme, purified from *H. volcanii* KM01-pJAM4351 grown on GlyMM, was used for activity assays. A. His-GK activity was assessed after a 1-hour incubation at 24°C, 33°C, 42°C, 51°C, 57°C, 60°C, and 65°C in 50 mM HEPES buffer, pH 8, containing 2 M NaCl. B. His-GK activity was assessed after a 1-hour incubation at the same temperatures in 50 mM HEPES buffer, pH 8, containing 0.1 M NaCl. C. His-GK activity was measured after incubation in 50 mM HEPES buffer, pH 8, with NaCl concentrations of 0.1, 0.15, 1, 2, 3, and 4 M. Samples were stored at 4°C, and enzymatic activity was measured at 0, 24, 48, and 72 hours. NA represents the control reaction, where His-GK was directly assayed from the freezer without incubation.

### Melting temperature of *H. volcanii* His-GK in high and low concentrations of glycerol

DSF was used to determine the apparent melting temperature (T_m_) and melting curve profiles of His- GK in the presence of low (1 %) and high (10 %) concentrations of glycerol (**Figure 8**). From this analysis, His-GK was found to display two distinct patterns of melt curves that were dependent on the glycerol concentration. In particular, the enzyme appeared more prone to interact with the fluorescent dye at moderate temperatures when assayed in the presence of high vs. low concentrations of glycerol **(Figure 8AB**). By contrast, the addition of glycerol at the two different concentrations had no significant impact on the apparent melting temperatures (T_m_ of 83-84 °C) of the enzyme calculated from the melt peaks of these profiles (**Figure 8CD**). The high T_m_ values identified by DSF were consistent with the extreme thermotolerance and resilience of this enzyme to harsh conditions. Moreover, the glycerol-dependent changes in melting curve profiles are consistent with the glycerol-induced shift in the equilibrium of the His-GK from dimer to a dimer-tetramer state, as observed by SEC.

**Figure 8.**
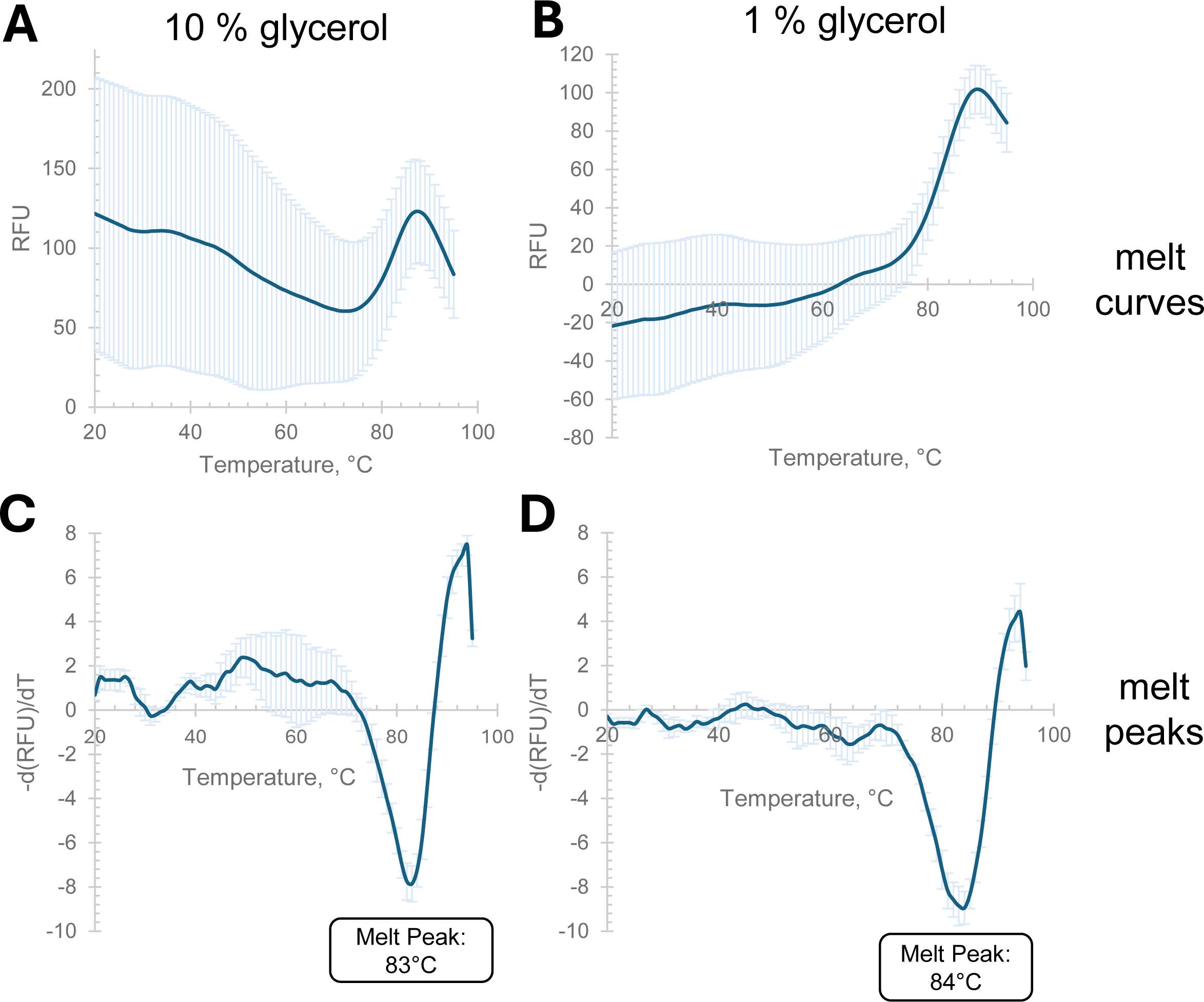
Thermal shift assay of *H. volcanii* glycerol kinase. His-GK, purified from *H. volcanii* KM01-pJAM4351 grown on GlyMM and stored at -80°C in 10% glycerol, was analyzed by thermal shift assay. The temperature was increased from 20°C to 95°C at a rate of 1°C per min. The top panels display the melt curves, while the bottom panels show the corresponding melt peaks. A and C: His-GK diluted in buffer containing 50 mM HEPES, pH 7.5, 2 M NaCl, and 10 % glycerol. B and D: His-GK diluted in the same buffer with 1 % glycerol. Comparison of melt peaks in the presence of 1 % vs. 10 % glycerol revealed no significant difference in T_m_ values (p- value = 0.08).

## DISCUSSION

### *H. volcanii* glycerol kinase displays biological function and can be purified to homogeneity when fused to an N-terminal poly-His tag

Affinity tags generated by fusion of a small polypeptide to the target protein are commonly used to enhance protein purification (32). Although vectors encoding these tags can be purchased commercially, choosing the right tag for optimal protein yield and identifying its best placement on the protein to maintain structure and function are still major challenges. The N-terminal poly-His-tag strategy has been used to successfully purify GKs from *E. coli*, *S. pneumoniae* and other organisms (33–39). However, the solubility and functionality of tagged proteins can vary. For instance, the N-terminal His-tagged GK from *M. pneumoniae* is insoluble, whereas the same protein with an N-terminal StrepII tag is soluble and functional (40).

In this study, the *H. volcanii* GK was evaluated for the effects of fusing a poly-His tag to its N-terminus compared to adding a StrepII tag to its C-terminus. Both GK variants were found to be biologically active and capable of supporting the growth of *H. volcanii* on glycerol as the sole carbon source. By contrast, significant differences were observed in purification yield between the two proteins with a 4.5-fold higher yield observed for His-GK compared to GK- StrepII. The 3 mg protein per L culture obtained for His-GK was useful for downstream biochemical analysis. Purification of GK with the His tag also resulted in a highly pure protein, as indicated by the absence of additional bands in SDS-PAGE gels. This type of yield and level of purity are robust when considering the host strain was *H. volcanii*. Using haloarchaeal hosts can be advantageous when working with their ‘salt-loving’ enzymes which often do not properly fold within or after recombinant expression in mesohalic hosts such as *E. coli* (41, 42).

Previous studies have shown that His-tagged proteins generally achieve higher purification efficiencies than StrepII-tagged proteins, although varying levels of purity have been reported in extracts from *E. coli*, yeast, *Drosophila*, and HeLa cells (43). Using high-salt buffers in our protocol likely contributed to reduced nonspecific interactions during His-tag purification (44). Furthermore, the binding of non-specific proteins observed in extracts from insect and mammalian cells due to high His residue content (45) was not an issue in our *H. volcanii* system as an improved strain (H1207) for expression of His-tagged proteins was used as the host (30). The positioning of the tag also influences purification yield and effectiveness. While both C-terminal and N-terminal tags have been used successfully, our 3D-modeling suggests that the N-terminal His tag provides an optimal balance between functionality and purification efficiency for *H. volcanii* GK. The results emphasize the importance of tailoring tag selection and positioning to the specific characteristics of the target protein.

### Optimal conditions in different glycerol kinases

Glycerol kinases from various organisms exhibit distinct optimal conditions that reflect their environmental adaptations. In *H. volcanii*, the GK appears optimized for extreme conditions and can accommodate a variety of metal cofactors including Mg^2+^, Mn^2+^ and Co^2+^. The enzyme’s catalytic activity operates over a wide pH range (6.0 to 10.0), is active in molar concentrations of salt, and functions at elevated temperatures, highlighting the adaptability of *H. volcanii* to hypersaline and thermophilic environments. Like the *H. volcanii* GK, many GKs use Mg^2+^ as the divalent metal cofactor (37, 46). However, most GKs display optimal activity at lower temperature, are more sensitive to salt, and prefer neutral to higher pH values. For example, the *T. brucei* GK is optimal at pH 7 to 9.5 (47), while the human GK is best suited for a neutral environment with optimal activity at pH 7.5 (37). Other archaea such as the hyperthermophilic *T. kodakarensis* provide additional insight into GKs with unusual biochemical properties, as this enzyme prefers pH 8.0 and 80 °C and favors cobalt ions (Co²⁺) over magnesium (Mg²⁺) for optimal activity (33). These findings emphasize that the choice of metal ion, pH, and other environmental factors significantly influence glycerol kinase activity across species. Enzymatic activity studies on *H. volcanii* GK further highlight how these environmental parameters — metal ion concentration, buffer type, temperature, pH, and salinity—shape its function.

### Effect of organic solvent and crude glycerol on *H. volcanii* GK activity

The organic solvent DMSO and crude glycerol, which contains methanol, were also examined for their effect on *H. volcanii* His-GK activity. These compounds have relevance to biotechnological applications in solvent-rich environments or crude feedstocks, where enzyme stability is essential. At 5 % DMSO, His-GK activity decreased with increasing salinity, but still retained about half of its activity when assayed with 5 % DMSO and 4 M NaCl compared to the no DMSO control at 0.1 M NaCl. This suggests that DMSO partially disrupts the enzyme’s activity at higher salinities. However, even at 10 % DMSO, His-GK maintained about 63 % of its activity across all salinity levels, indicating resilience to relatively high concentrations of DMSO in high-salinity conditions. Many halophilic enzymes are well-known for their ability to perform in salt- and solvent-rich environments, which are common in industrial processes. For instance, *H. volcanii* alcohol dehydrogenase (ADH2) shows remarkable stability in presence of DMSO and methanol (48), and *H. volcanii* laccase (LccA) retains over 50 % of its activity after 24 h in DMSO (49). Similarly, the NAD^+^-dependent glutamate dehydrogenase from *Halobacterium salinarum* exhibits solvent tolerance (50). Moreover, His-GK demonstrated equivalent efficiency in degrading crude glycerol as it did with pure glycerol. This suggests that the enzyme is not only tolerant to methanol but also capable of efficiently catalyzing glycerol degradation in complex feedstocks. This property enhances His-GK’s potential for applications in biodiesel production and glycerol waste bioremediation.

### Stability of *H. volcanii* His-GK activity at varying temperatures and salinity levels

The tolerance of *H. volcanii* His-GK to harsh environmental conditions was also evaluated. At 2 M NaCl, the enzyme retained full activity across all tested temperatures (24°C to 65°C), showcasing its thermal stability under high-salinity conditions. In contrast, at 0.1 M NaCl, His- GK was stable only up to 51 °C, with significant activity loss at 57 °C and complete denaturation at 60 °C and 65 °C. These results underscore the critical role of salinity in preserving His-GK’s structural integrity and enzymatic activity at elevated temperatures. Interestingly, the thermal stability of *S. cerevisiae* mesophilic GK retains full activity up to 50°C after 1 h incubation and exhibits activity loss beyond this threshold (51).

### Impact of glycerol on His-GK stability

Thermal shift assays revealed that the *H. volcanii* GK has a high melting temperature and suggested that glycerol induced an alteration to the enzyme structure. Irrespective of glycerol, the T_m_ of His-GK was found to be similar at 83 to 84 °C. However, the conformational heterogeneity of the melt profile observed at 10 % vs. 1 % glycerol suggests dynamic structural transitions. Similar trends in high T_m_ values have been observed in other GKs including the thermophilic fungus *C. thermophilum* GK, where the T_m_ is 60 °C (38). Interestingly, the T_m_ of *C. thermophilum* GK is reduced to 53-55 °C in the presence of glycerol, ATP, or ADP, while increased when both nucleotide and glycerol are included in the assay, with T_m_ values reaching 63.6 °C for the ATP-glycerol complex and 63 °C for the ADP-glycerol complex (38). While the His-GK showed only minor T_m_ shifts with glycerol, the observed structural dynamics in the melding profiles may reflect a glycerol-induced mechanism analogous to *C. thermophilum* GK, where ligand interactions enhance enzyme performance (38).

### *H. volcanii* glycerol kinase exhibits a glycerol-dependent conformational shift

Here we find His-GK shifts from a dimer to a dimer-tetramer equilibrium in the presence of glycerol. This shift mirrors patterns observed in GKs from other organisms. For example, the *E. coli* GK exhibits a dimer-tetramer equilibrium in the presence of FBP, with the dimer being the active form and the tetramer the inactive form (52). Likewise, the *Enterococcus casseliflavus* GK exhibits an active dimer conformation but forms a pseudo-tetramer in specific conditions (53). By comparison, GKs from *Plasmodium falciparum*, *Trypanosoma brucei gambiense*, and the thermophilic fungus *Chaetomium thermophilum* maintain active dimeric conformations (38, 54, 55). Interestingly, the archaeal *T. kodakarensis* GK shifts from a dimer to a hexamer in the presence of glycerol. This conformational shift results in a closed structure that enhances ATP affinity tenfold, effectively regulating enzymatic activity (56).

### Positive cooperativity and kinetic parameters of *H. volcanii* glycerol kinase

The glycerol- induced shift in the oligomeric state corresponds with our finding that *H. volcanii* GK exhibits sigmoidal kinetics and positive cooperativity for glycerol, ATP, and magnesium, as determined by the Hill equation. This finding suggests this enzyme is adapted to rapidly respond to variations in substrate levels in a positive manner. This dynamic behavior aligns with the organism’s adaptation to fluctuating environmental conditions and preference for glycerol over glucose. The Hill coefficients are strong indicators of cooperativity for ATP (n = 2.35 ± 0.080) and glycerol (n = 2.12 ± 0.188) with the magnesium cofactor (n = 1.6 ± 0.15) suggested to have some level of cooperative interaction, but less pronounced compared to the other effectors. By contrast, magnesium showed the strongest binding affinity to the enzyme with a dissociation constant (*K*_d_) of 89 ± 6 µM and a catalytic efficiency (*k*_cat_/*K*_d_) of 1,710 ± 150 mM⁻¹ s⁻¹. The high sensitivity of *H. volcanii* GK to magnesium is consistent with other GKs, which are used in as biocatalysts in clinical and industrial settings, such as in the colorimetric detection of magnesium levels in serum (57).

The *H. volcanii* GK exhibits kinetic properties for ATP and glycerol that are distinct compared to other GKs. The relatively high *K*_d_ values for glycerol (1.44 ± 0.16 mM) and ATP (1.6 ± 0.08 mM) indicate that *H. volcanii* GK has a low affinity for these substrates compared to its bacterial and eukaryotic counterparts. For example, the gram-positive bacterium *P. pentosaceus* GK has *K*_m_ values of 0.11 mM and 0.37 mM for glycerol and ATP, respectively (46). *E. coli* GK has a *K*_m_ of 0.01 mM for glycerol (58). Eukaryotic GKs exhibit *K*_m_ values for ATP and glycerol in the range of 0.002 to 0.055 mM. Specifically, the *K*_m_ values for glycerol and ATP are 0.01 mM in rat liver, 0.02 and 0.01 mM in beef liver, 0.002–0.003 and 0.01 mM in human liver, and 0.035 and 0.055 mM in *Candida mycoderma*, respectively (59).

Interestingly, *H. volcanii* GK’s kinetic properties also differ from the archaeal *T. kodakarensis* GK, which demonstrates higher affinity for ATP (*K*_m_ of 0.0154 mM) and glycerol (*K*_m_ of 0.111 mM) (33). The ATP-binding affinity of the *H. volcanii* GK also contrasts with the GK of the thermophilic fungus *C. thermophilum*, where substrate inhibition occurs beyond 0.4 mM ATP concentrations (38). These variations reflect the evolutionary divergence and environmental adaptations of GKs across domains of life. These also findings highlight the distinct metabolic requirements and habitats of halophilic archaea which prefer glycerol and have an excess supply of this carbon source in hypersaline environments.

The positive cooperativity of *H. volcanii* GK for glycerol, ATP and magnesium, has not been observed for other GKs. Positive cooperativity is often associated with allosteric regulation of multisubunit enzymes structures. This kinetic behavior aligns with our finding that *H. volcanii* GK undergoes a glycerol-induced transition from a dimer to a dimer-tetramer equilibrium, which may enhance substrate affinity at other active sites. While this type of behavior has not been identified in other GKs, allosteric regulation is a phenomenon well-documented in GKs. For example, the *E. coli* GK is inhibited by the allosteric effectors FBP and EIIA^Glc^, where FBP promotes a conformational change in the enzyme from a functional dimeric state to a less active tetrameric state (52, 60). In this mechanism, the bacterial GK exhibits “positive cooperativity”, meaning that the binding of one FBP molecule to the enzyme increases the affinity for subsequent FBP molecules to bind, leading to a more significant inhibition effect at lower FBP concentrations. Moreover, the *E. coli* GK displays negative cooperativity in ATP binding, apparently helping to stabilize the ATP-Mg²⁺ complex, prevent excessive ATP occupancy, avoid overactivity of the enzyme (61). The glucose-specific enzyme IIA (EIIA^Glc^) of the phosphotransferase system (PTS) is also an allosteric inhibitor of the *E. coli* GK (60). Together these mechanisms of allosteric regulation of the E. coli GK provide a control mechanism to facilitate diauxic growth on glucose over glycerol and ensure metabolic balance with FBP the dominant allosteric control mechanism (62). FBP does not appear to be a physiological regulator of eukaryotic GKs where it causes a slight if any inhibitory effect on GKs from organisms such as the protozoan parasite *P. falciparum* (55), the thermophilic fungus *C. thermophilum* (38) and the salt-tolerant yeast *Debaryomyces hansenii* (63). Likewise, FBP is not an allosteric inhibitor of the *H. volcanii* GK, which is consistent with the organism’s preference of glycerol over glucose as a carbon source (7).

The biochemical properties of the *H. volcanii* GK indicate its suitability for industrial applications. The enzyme’s ability to maintain robust biocatalytic activity across a broad spectrum of environmental conditions underscores its durability, particularly in applications where factors such as salinity, pH, organic solvents, and temperature can fluctuate. Its distinct substrate affinities and cooperativity characteristics offer chances for customizing biocatalytic applications via protein engineering, paving the way for progress in sustainable biotechnology and bioeconomic efforts. Comparative analysis with GKs exhibiting low substrate affinities, such as *Trypanosoma brucei* GK, where alanine-to-serine mutations enhance glycerol affinity (36), may further elucidate the molecular factors that influence substrate specificity and efficiency in *H. volcanii* GK. These findings could guide the development of optimized enzymes for industrial glycerol applications, promoting innovative approaches in bioprocessing. The capacity of *H. volcanii* GK to maintain activity in extreme conditions, along with its cooperative interaction with glycerol, ATP, and magnesium, positions it as a compelling option for biotechnological applications. Additional research into the structural mechanisms that promote positive cooperativity of the *H. volcanii* GK will be advantageous for enhancing the effectiveness of GKs in industrial applications, such as the bioconversion of waste glycerol into value added products.

## Conclusion

The positive cooperativity and kinetic properties of *H. volcanii* GK emphasize its potential for use as a robust enzyme in biotechnological processes under extreme conditions. The enzyme’s cooperative kinetics, structural flexibility, and efficient substrate utilization distinguish it from GKs found in other organisms, providing valuable insights into the design of industrial processes that utilize extremophilic enzymes. Further exploration of the enzyme’s structural and regulatory mechanisms will deepen our comprehension of its functionality and broaden its applications in industrial biotechnology.

## Acknowledgments

Funds awarded to JMF to characterize archaeal biocatalysts and fundamental biological mechanisms were through the U.S. Department of Energy, Office of Basic Energy Sciences, Division of Chemical Sciences, Geosciences and Biosciences, Physical Biosciences Program (DOE DE-FG02-05ER15650) and National Institutes of Health (NIH R01 GM57498), respectively.

